# Glioblastoma cells taste temozolomide via TAS2R43

**DOI:** 10.1101/2025.06.27.661979

**Authors:** Ana R. Costa, Ana C. Duarte, Isabel Gonçalves, Robert Preissner, José F. Cascalheira, Helena Marcelino, Cecília R.A. Santos

**Author notes:** Corresponding author: Dr Cecília R. A. Santos, RISE-Health, UBI, Faculty of Health Sciences, University of Beira Interior, Av. Infante D. Henrique, 6200-506 Covilhã, Portugal.

## Abstract

Bitter taste receptors (TAS2Rs) have a widespread expression in various extraoral organs where they detect the chemical composition of body fluids and trigger biological responses accordingly. Among the chemicals recognised by TAS2Rs there are natural and synthetic compounds including therapeutic drugs. We have shown that TAS2Rs are expressed in the blood-cerebrospinal fluid barrier where they regulate efflux transporters, thereby controlling the transport of compounds into the cerebrospinal fluid. More recently, we assessed the expression of these receptors in human glioblastoma cells, where 20 out of the 26 human TAS2Rs were identified. In this study, we investigated if temozolomide, the standard chemotherapy for glioblastoma, activates the bitter signalling pathway with impact in its therapeutic efficacy. Notably, we found that blocking the bitter taste signalling pathway significantly reduced the anti-proliferative and pro-apoptotic effects of temozolomide, and identified TAS2R43 as the receptor mediating these effects. We propose that upon ligand binding, TAS2R43 modulates multidrug resistance proteins (MDRs) activity, facilitating temozolomide entrance into glioblastoma cells. These findings underscore the importance of the taste transduction pathway in evaluating the chemical composition of the glioblastoma microenvironment. Furthermore, our data suggest that TAS2R43 could serve as a biomarker for the efficacy of temozolomide and other drugs that are substrates of MDRs.

## 1. Introduction

Apart from sensing bitter compounds in the oral cavity, TAS2Rs were also described in several extraoral tissues, like the gastrointestinal tract, airway epithelium, heart, testis and brain (Avau & Depoortere 2016; Duarte et al. 2020a; Jaggupilli et al. 2017; Li 2013). Experimental and clinical data revealed that TAS2Rs, which belong to the large family of G protein-coupled receptors, and their ligands, have a crucial role in cancer progression and metastasis, by controlling many aspects of tumorigenesis, proliferation, migration, cancer cell invasion and cancer-related signalling pathways (Dorsam & Gutkind 2007; O’Hayre et al. 2014). Despite their importance as therapeutic targets in cancer treatment, they have rarely been exploited in cancer except for certain endocrine and hormone-responsive tumours (Lappano & Maggiolini 2011; Usman et al. 2020).

Glioblastoma is the most common and aggressive form of primary brain cancer classified grade 4 by the World Health Organization, with a 5-year survival of approximately 3% and a median survival of less than 18 months (Louis et al. 2021; Reni et al. 2017). The current golden standard for glioblastoma treatment is palliative and includes surgery, radiotherapy and temozolomide (TMZ) chemotherapy. TMZ is an alkylating agent known for its ability to alkylate/methylate DNA, resulting in DNA damage and tumour cells death by apoptosis (Strobel et al. 2019). However, its efficacy is limited because some tumour cells can repair TMZ-induced DNA damage through the action of O6-methylguanine DNA methyltransferase gene (Kaina et al. 2007), relapsing of chemo- and radioresistant cancer stem-like cells and because these are highly infiltrative tumours (Bao et al. 2006; Chen et al. 2012; Eramo et al. 2006). Additionally, brain barriers, and brain cancer cells (Gomez-Zepeda et al. 2020; Hermann et al. 2006; Qosa et al. 2015; Saunders et al. 2016), overexpressed multidrug efflux transporters which extrude several CNS-targeting medicines (Gomez-Zepeda et al. 2020; Gupta et al. 2020; Hermann et al. 2006; Qosa et al. 2015; Saunders et al. 2016), which fail to achieve therapeutic concentrations at their target cells (Pitcher & Quevedo 2016). However, little is known about the mechanisms underlying their regulation. We recently identified 20 out of the 26 human TAS2R in glioblastoma cells that mediate signal transduction in response to a wide variety of bitter agonists (Adler et al. 2000; Chandrashekar et al. 2000; Lang et al. 2023).

We hypothesized that these TAS2Rs which are differentially expressed in astrocytes *and* glioblastoma cells, may bind TMZ, given its bitter properties. We first analysed if TMZ activates the bitter taste signalling pathway in glioblastoma cells and found that the presence of TAS2R43 is important for its anti-cancer efficacy in these glioblastoma cell lines.

## 2. Materials and methods

### 2.1 Materials

Temozolomide (TMZ; CAS No 85622-93-1) was purchased from Cayman Chemical (#14163). A stock solution was prepared in dimethyl sulfoxide (DMSO) and freshly dissolved in Tyrode’s solution or culture medium before the experiments, where the DMSO final concentration did not exceed 1%. A vehicle control was included in all the experiments. Probenecid (CAS No 57-66-9), a known bitter taste receptors’ antagonist, was obtained from Sigma-Aldrich (#P8761), dissolved in 1 N NaOH at 0.17 M and diluted in Tyrode’s solution or culture medium. FURA-2AM (#F1221), pluronic acid F-127, Lipofectamine™ 2000 (#11668027), Opti-MEM medium (#11058-021), small interfering RNA (siRNA) targeting α-gustducin (GNAT3; #4392420; ID s51191; sequence: S-*GCGAGAUGCAAGAACCGUATtt* and AS-*UACGGUUCUUGCAUCUCGCtc*) and bitter taste receptor TAS2R43 (#AM16708; ID 202331; sequence: S-*CCCUACUAUCUUUUAUGCUtt* and AS-*AGCAUAAAAGAUAGUAGGGtc*), and scramble siRNA (#4390843) were purchased in ThermoFisher Scientific. MTT [3-(4,5-dimethylthiazol-2-yl)-2,5-diphenyltetrazolium bromide] was purchased from Gerbu Biotechnik GmbH (#1006).

### 2.2 Cell Culture

Human malignant glioblastoma cell lines U-87MG and U-373MG were kindly provided by Dr. Joseph Costello (University of California, San Francisco), and SNB-19 was obtained from German Collection of Microorganisms and Cell Cultures. Glioblastoma cells were grown in Dulbecco’s modified Eagle’s medium (DMEM) high glucose with stable glutamine (bioWest #L0103) supplemented with 10% (v/v) FBS and penicillin (100 IU/mL)/streptomycin (100 μg/mL), and incubated in a humidified atmosphere containing 5% CO_2_ at 37 °C.

### 2.3 Evaluation of the impact of TAS2R in TMZ cytotoxicity

#### 2.3.1 Effects of TMZ in intracellular Ca^2+^ responses of glioblastoma cells

The effects of TMZ on Ca^2+^ cells responses, in the presence or absence of the TAS2Rs antagonist probenecid, were assessed by single-cell Ca^2+^ imaging assays in glioblastoma SNB-19 and U-373MG cells. Briefly, approximately 3.5x10^4^ glioblastoma cells were seeded in μ-slide 8 well ibiTreat (Ibidi # 80826) and grown until 60-70% confluency, followed by measurement of changes in intracellular Ca^2+^ levels after stimulation. Glioblastoma cells were loaded with 5 μM of FURA-2 AM and 0.02% pluronic acid F-127 in culture medium for 45 min. Next, cells were washed twice with Tyrode’s solution pH 7.4 [NaCl 140 mM, KCl 5 mM, MgCl_2_ 1.0 mM, CaCl_2_ 2.0 mM, Na-pyruvate 10 mM, Glucose 10 mM, HEPES 10 mM, NaHCO_3_ 5.0 mM] and loaded with Tyrode’s for 30 min. After that, dose-response experiments were performed with a range of TMZ concentrations (50-200 µM), in the presence or absence of 30 min incubation with probenecid (1 mM). The μ-slide plates were placed on a Widefield Axio Observer Z1 inverted microscope (Zeiss). Stock solution of TMZ and probenecid was freshly prepared in Tyrode’s solution before the experiments. The stimulus was applied manually with a micropipette after baseline was recorded. The intracellular Ca^2+^ levels were evaluated by quantifying the ratio of the fluorescence emitted at 520 nm following alternate excitation at 340 nm and 380 nm, using a Lambda DG4 apparatus (Sutter Instrument) and a 520 nm bandpass filter (Zeiss) under a Fluar 40x/1.30 Oil M27 objective (Zeiss). Data were processed using the Fiji software (MediaWiki). Changes in fluorescence ratio (F=F340/F380) were measured in at least 20 cells, in three or more independent experiments. Response intensity, or intracellular Ca^2+^ variation (ΔF/F0), was calculated in the following way: ΔF/F0=(F-F0)/F0, where F0 corresponds to fluorescence ratio average at baseline (2 min acquisition before stimulus) and F correspond to maximum peak of fluorescence ratio evoked by the stimulus applied to the cells.

#### 2.3.2 Evaluation of the impact of GNAT3 knockdown in TMZ cytotoxicity

After Ca^2+^ imaging experiments, the cytotoxicity of TMZ was assessed in glioblastoma cells in the presence or absence of the TAS2Rs antagonist probenecid, using the MTT assay. Additionally, the specific activation of bitter taste signalling by TMZ was assessed in glioblastoma cells after GNAT3 knockdown with specific siRNA. Briefly, glioblastoma cells were grown until 60% confluency or transfected for 24 h with a mixture of GNAT3 siRNA (10 nM) and Lipofectamine™ 2000 in Opti-MEM medium, following the manufacturer’s instructions. A scramble siRNA (10 nM) was also used as negative control for GNAT3 specific targeting. Then, cells were incubated for 72 h with TMZ (500 μM) or vehicle (DMSO 1%), in the presence or absence of 1 mM probenecid diluted in culture medium. Then, 100 μL culture medium was removed and 10 μL of MTT solution (5 mg/mL in PBS) was added for approximately 45 min at 37 °C in a humidified atmosphere containing 5% CO_2_. Untreated cells and ethanol 70% treated cells were used as negative and positive controls, respectively. Following MTT incubation, formazan crystals were dissolved in DMSO for 15 min, and absorbance was read at 570 nm in a microplate spectrophotometer xMark™ (Bio-Rad). The viability of glioblastoma cells was expressed as a percentage of the absorbance determined in the vehicle control. The apoptotic effect of TMZ, in the presence or absence of probenecid, was also evaluated in glioblastoma cells by Hoechst 33342 nuclei staining. Glioblastoma cells were seeded in a coverslip and the stimuli were carried out by incubating the cells for 72 h with TMZ (500 μM) or vehicle (DMSO 1%), in the presence or absence of 1 mM probenecid diluted in culture medium. After removing the medium, cells were fixed with PFA 4% for 10 min followed by incubation for 10 min with Hoechst 33342 (diluted 1:1000). After several washes, cells were mounted onto microscope slides and visualized under a confocal microscope LSM 710 (Zeiss) using a magnification of 63x (Plan-Apochromat 63x/1.4 Oil DIC M27). Apoptotic cells were distinguished from healthy or necrotic cells by the observation of condensed DNA and fragmented nuclei. The apoptotic rate was calculated using the ratio between the number of apoptotic cells and the total number of cells.

### 2.4 Analysis of the role of TAS2R43 on the glioblastoma cells’ response to TMZ

Following the demonstration that the bitter taste signalling pathway was involved in the mediation of the effects of TMZ, we assessed *in-silico* which TAS2R could have a higher likelihood of binding to TMZ according to the webserver VirtualTaste method (Fritz et al. 2021). Using this approach, six TAS2Rs, namely TAS2R38 and TAS2R43, were identified as possible target-receptors for TMZ. Both had been detected in glioblastoma cells (U-87MG, SNB-19, and U-373MG) by RT-PCR. Additionally, TAS2R43 protein detection in glioblastoma cells was assessed by immunocytochemistry, as described before (Costa et al. 2025 - bioRxiv submission ID BIORXIV/2025/661264). Finally, the role of TAS2R43 on the response to TMZ was assessed in glioblastoma cells after TAS2R43 knockdown with specific siRNA, as described in section 2.3.2 for GNAT3.

### 2.5 Data Analysis

Statistical analysis and dataset comparisons were performed using GraphPad Prism 9.3.1 (GraphPad Software). Statistical significance was determined by One-Way ANOVA followed by the software’s recommended multiple comparisons post-hoc test. Results are presented as mean ± SEM of at least three independent experiments, and data were considered statistically different for a p-value<0.05.

## 3. Results

### 3.1 TMZ elicits Ca^2+^ uptake in glioblastoma cell lines

We assessed SNB-19 and U-373MG response to TMZ (10–200 μM) using a Ca^2+^ functional assay (Figure 1) in the presence or absence of probenecid, a blocker of TAS2R16, R38, and R43, to elucidate the role of TAS2Rs in the response of glioblastoma cells to TMZ (Greene et al. 2011; Wölfle et al. 2015). Our results showed that TMZ was able to trigger a functional response in glioblastoma cells. TMZ at 100 µM (ΔF/F0=0.220±0.039) triggered a significant increase in intracellular Ca^2+^ levels in SNB-19 cells (Figure 1A) in comparison to the vehicle (ΔF/F0=0.025±0.008), abolished in the presence of probenecid (ΔF/F0=0.025±0.006). U-373MG cells stimulated with 50 and 100 µM TMZ showed higher Ca^2+^ levels (ΔF/F0=0.437±0.137 and ΔF/F0=0.718±0.155) in comparison with vehicle control (ΔF/F0=0.052±0.007) (Figure 1B). In addition, the TMZ effects on intracellular Ca^2+^ levels in the presence of probenecid were reverted (ΔF/F0=0.034±0.012). Neither 10 µM nor 200 µM TMZ triggered significant Ca^2+^ responses in both SNB-19 and U-373MG glioblastoma cells (Figure 1).

**Figure 1.**
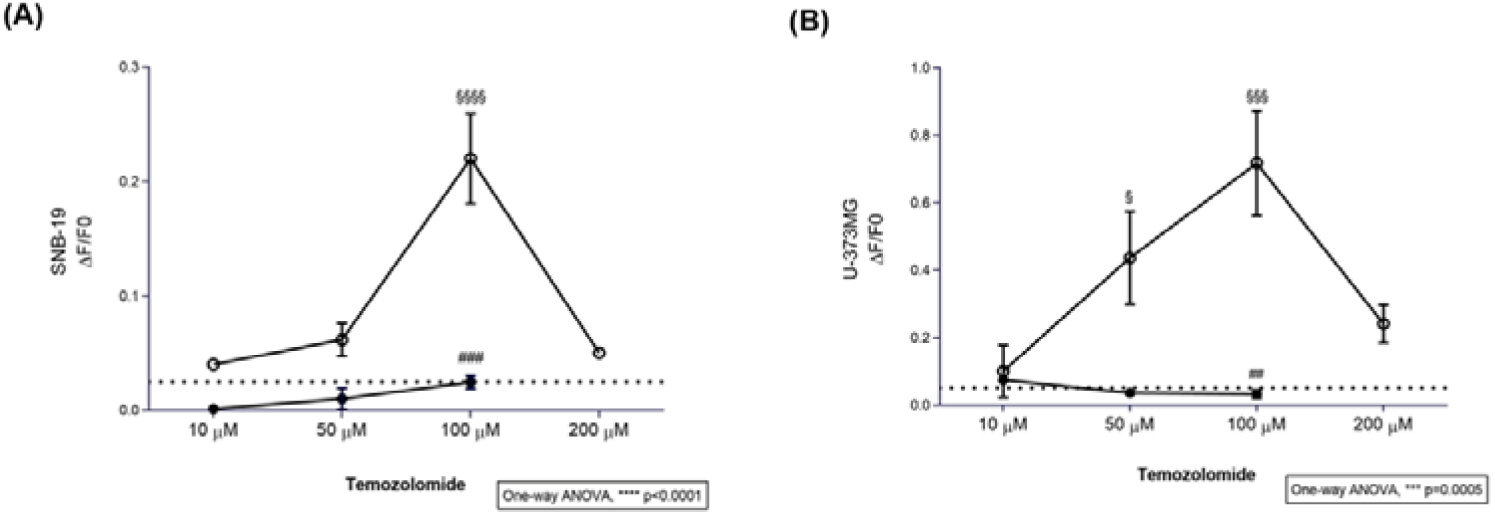
Calcium dose-response curves of SNB-19 and U-373MG glioblastoma cells to temozolomide. Different concentrations of temozolomide (TMZ; 10-200 μM), in the presence (●) or absence (○) of 1 mM probenecid, a known TAS2R inhibitor. Dot line: Ca^2+^ levels measured in cells with vehicle only (DMSO ≤ 0.2%). Results are presented as mean ± SEM. Statistical analysis was performed by two-way ANOVA followed by Tukey’s multiple comparisons test. [N≥3 independent experiments; ^§^versus vehicle; ^#^versus probenecid+TMZ].

### 3.2 The bitter taste signalling pathway enhances the anti-cancer effects of TMZ

Next, we proceeded to dose-response (50-500 µM) cytotoxicity assays, and found that 500 µM TMZ reduced the viability of U-87MG (49.08%±1.92), SNB-19 (44.46%±0.97) and U-373MG (50.92%±0.45) cells (Figure 2A).

**Figure 2.**
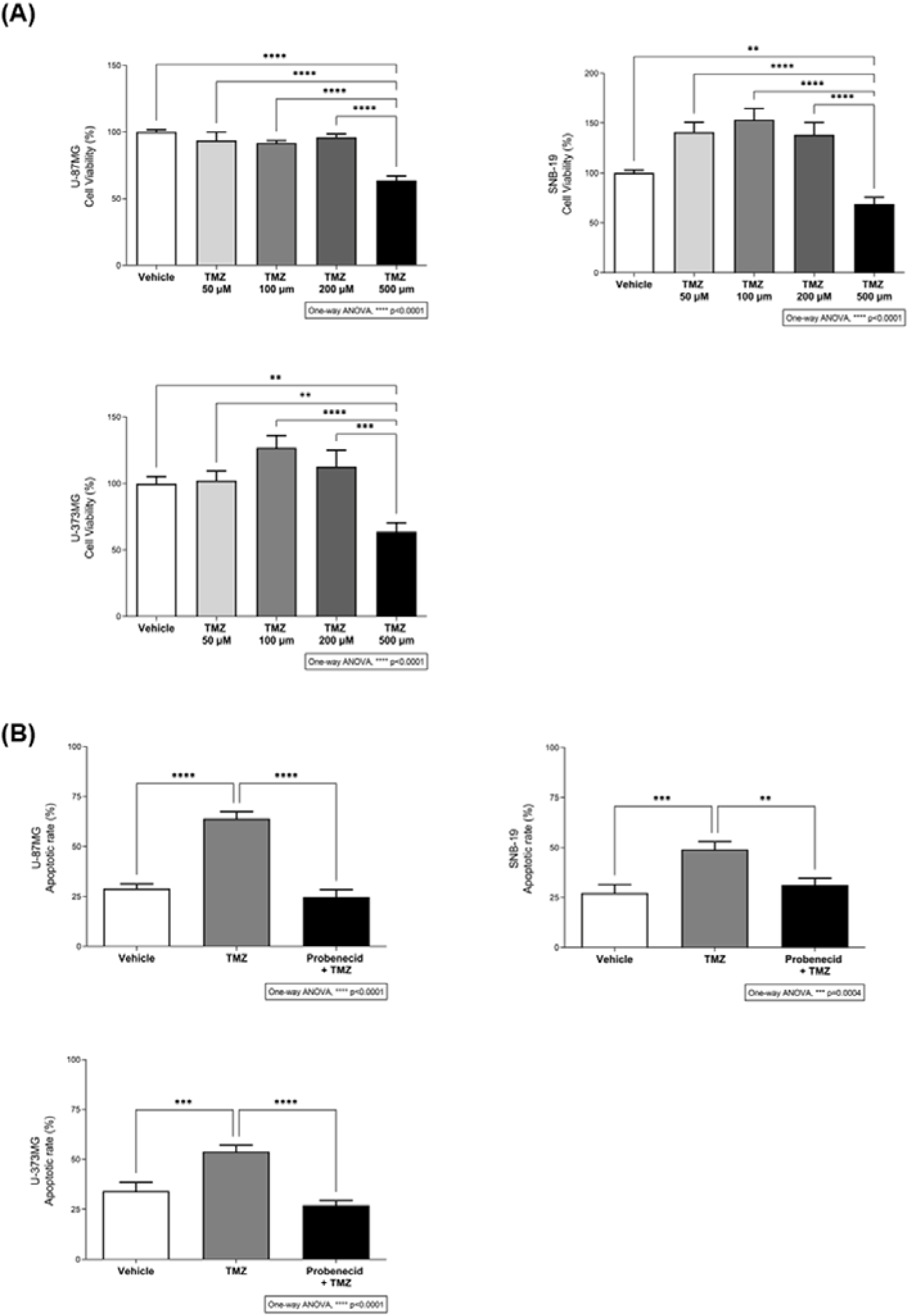
Cytotoxic effect of different concentrations of temozolomide. (A) Dose-response cytotoxicity assay of glioblastoma cells exposed to different concentrations of temozolomide (TMZ; 50-500 μM) for 72 hours was assessed in U-87MG, SNB-19 and U-373MG glioblastoma cells by MTT assay. Results are presented as mean ± SEM. Statistical analysis was performed by one-way ANOVA followed by Tukey’s multiple comparisons test. [N≥3 independent experiments; **p<0.01, ***p<0.001 and ****p<0.0001]. (B) The apoptotic rate of U-87MG, SNB-19 and U-373MG glioblastoma cells incubated with 500 µM TMZ for 72h, in the presence or absence of 1 mM probenecid, was carried out by Hoechst 33342 staining (data not shown). Bar graphs represent mean ± SEM. Statistical analysis was performed by one-way ANOVA followed by Tukey’s multiple comparisons test. [N≥3 independent experiments; **p<0.01, ***p<0.001 and ****p<0.0001].

Then, we evaluated the effects of TMZ on the apoptosis of glioblastoma cells in the presence of probenecid, a known TAS2R16, R38, and R43 inhibitor (Greene et al. 2011), by staining the cell nuclei with Hoechst 33342 and counting the nuclei with apoptotic vesicles. As expected, TMZ induced apoptosis in these cells, but this effect was reduced in the presence of probenecid (Figure 2B). Our observations put in evidence that TMZ may be a TAS2Rs ligand and that the taste signalling pathways modulates its effects or penetration in these cells. To test this hypothesis, we proceeded with knockdown experiments, of the specific guanine nucleotide binding protein that mediates taste receptor mediated signalling cascades, GNAT3 (Figure 3). TMZ alone induced a reduction in the cells’ viability of approximately 37.4% in U-87MG, 41.1% in SNB-19, and 45.9% in U-373MG in comparison with non-, mock- and siRNA scramble-transfected cells (Figure 3B). Notably, the presence of probenecid, or the silencing of GNAT3, reduced the effects of TMZ on the viability of glioblastoma cells (Figure 3B), reinforcing the results obtained in the previous experiments. Overall, TMZ elicited Ca^2+^ responses in a dose-dependent manner and showed cytotoxic effects in glioblastoma cells, which were significantly reduced in the presence of probenecid or by GNAT3 silencing, highlighting that TMZ effects in these cells depend on the activation of the bitter taste signalling pathway.

**Figure 3.**
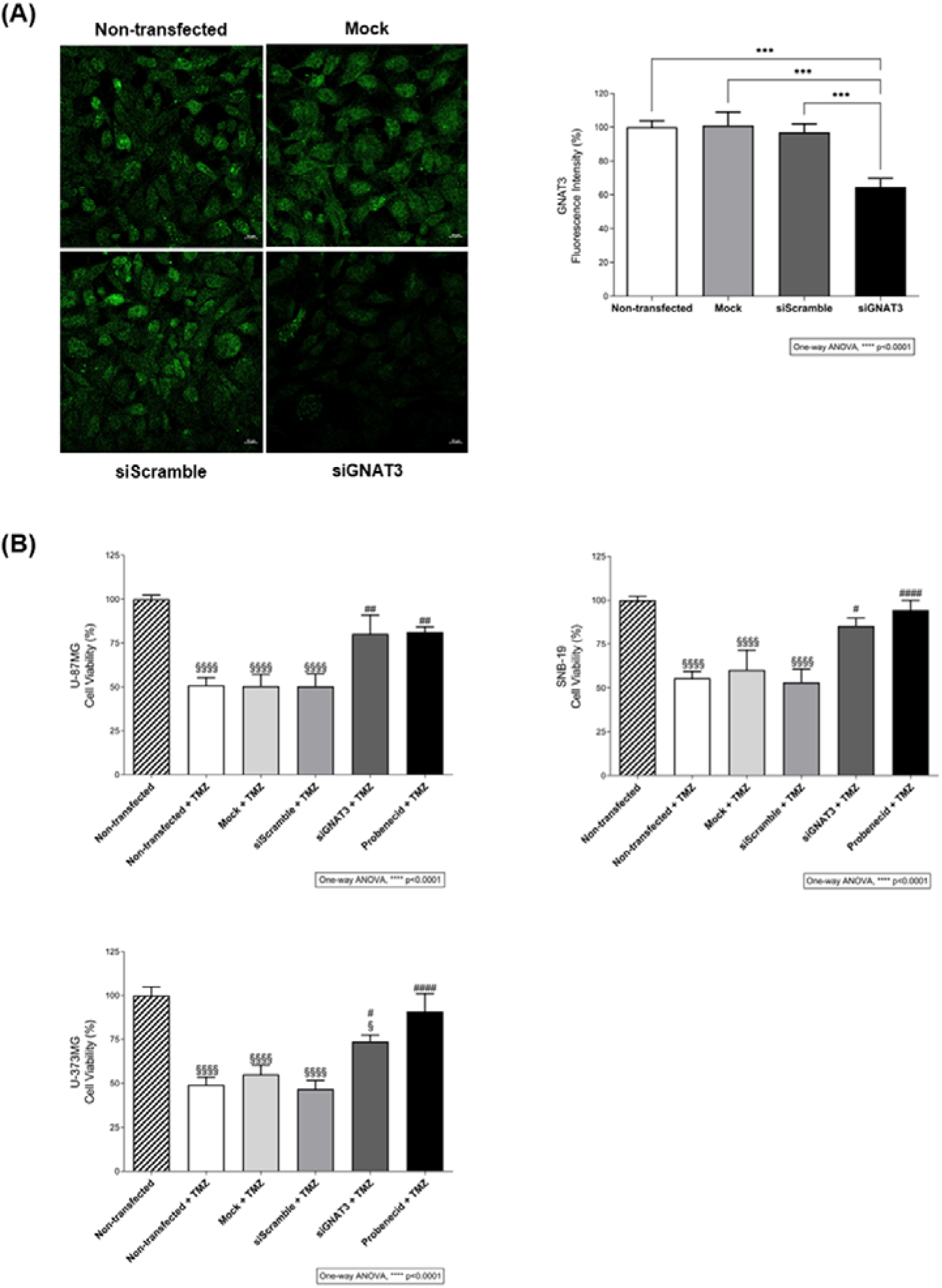
The effect of temozolomide on the viability of glioblastoma cells requires the activation of the bitter taste signalling pathway. (A) Immunofluorescence analysis of α-gustducin (GNAT3) expression after siRNA transfection in SNB-19 glioblastoma cells. Protein levels of siRNA GNAT3-transfected cells are decreased in comparison with non-, mock-, and siRNA scramble-transfected cells. The quantification of GNAT3 fluorescence intensity (green) was performed in different regions of interest of confocal microscopy images obtained from three independent experiments. Nuclei were stained with Hoechst 33342 (not shown). Scale bar: 10 µm. Bar graphs represent mean ± SEM. Statistical analysis was performed by one-way ANOVA followed by Tukey’s multiple comparisons test. [N=3 independent experiments; ***p<0.001]. (B) Effects of TMZ in the viability of U-87MG, SNB-19 and U-373MG glioblastoma cells transfected or mock-transfected for 24 h with GNAT3 or a scramble siRNA, and incubated with 500 µM TMZ for 72 h, in the presence or absence of 1 mM probenecid. Bar graphs represent mean ± SEM. Statistical analysis was performed by one-way ANOVA followed by Tukey’s multiple comparisons test. [N≥3 independent experiments; *p<0.05, **p<0.01 and ****p<0.0001; ^§^versus non-transfected; ^#^versus non-transfected+TMZ].

### 3.3 TAS2R43 mediates the effects of TMZ in the viability of glioblastoma cells

After confirming that the bitter taste signalling pathway has a role in the response of glioblastoma cells to TMZ, the next step was identifying which TAS2R(s) TMZ binds to. Based on the webserver VirtualTaste predictions (Figure 4A), six TAS2Rs could recognize TMZ: R38 (0.72) > R10 (0.68) > R43 (0.65) > R45 (0.62) > R44 (0.60) > R14 (0.59). However, only TAS2R38 and R43 bind probenecid [27].

**Figure 4.**
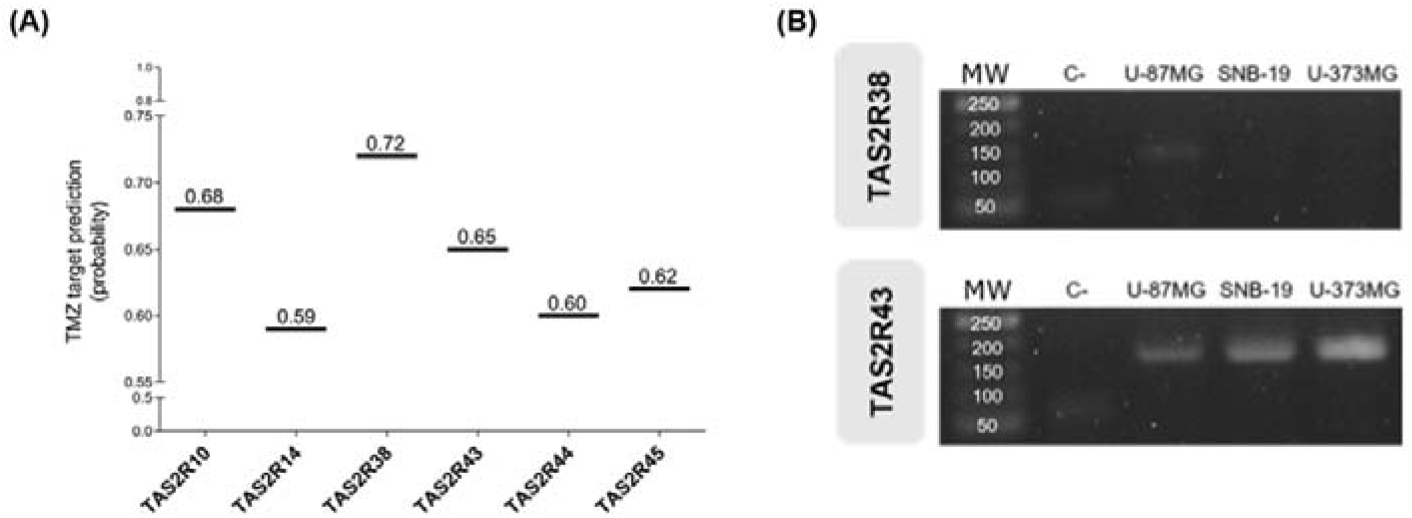
TMZ target prediction. (A) TMZ is predicted to bind TAS2R38 > R10 > R43 > R45 > R44 > R14 according to the webserver VirtualTaste algorithm. Results are presented as probability. (B) mRNA expression profile of TAS2R38 and R43 in glioblastoma cell lines (U-87MG, SNB-19, U-373MG). Only TAS2R43 mRNA and protein were detected in all cell lines. The identities of the amplified products were confirmed by Sanger sequencing. MW: molecular weight (base pair); C-: negative control.

Therefore, as only TAS2R43 was expressed in all the glioblastoma cell lines analysed (Figure 4B and 4C) we proceeded with the analysis of glioblastoma cells viability upon TAS2R43 knockdown (Figure 5A). TMZ alone induced a reduction of approximately 41.78±2.44% in the viability of U-87MG, SNB-19 and U-373MG cells in comparison with non-, mock- and siRNA scramble-transfected cells but was unable to reduce the viability of TAS2R43-silenced cells (Figure 5B), suggesting that the reduction of cell viability induced by TMZ depends on TAS2R43 activation.

**Figure 5.**
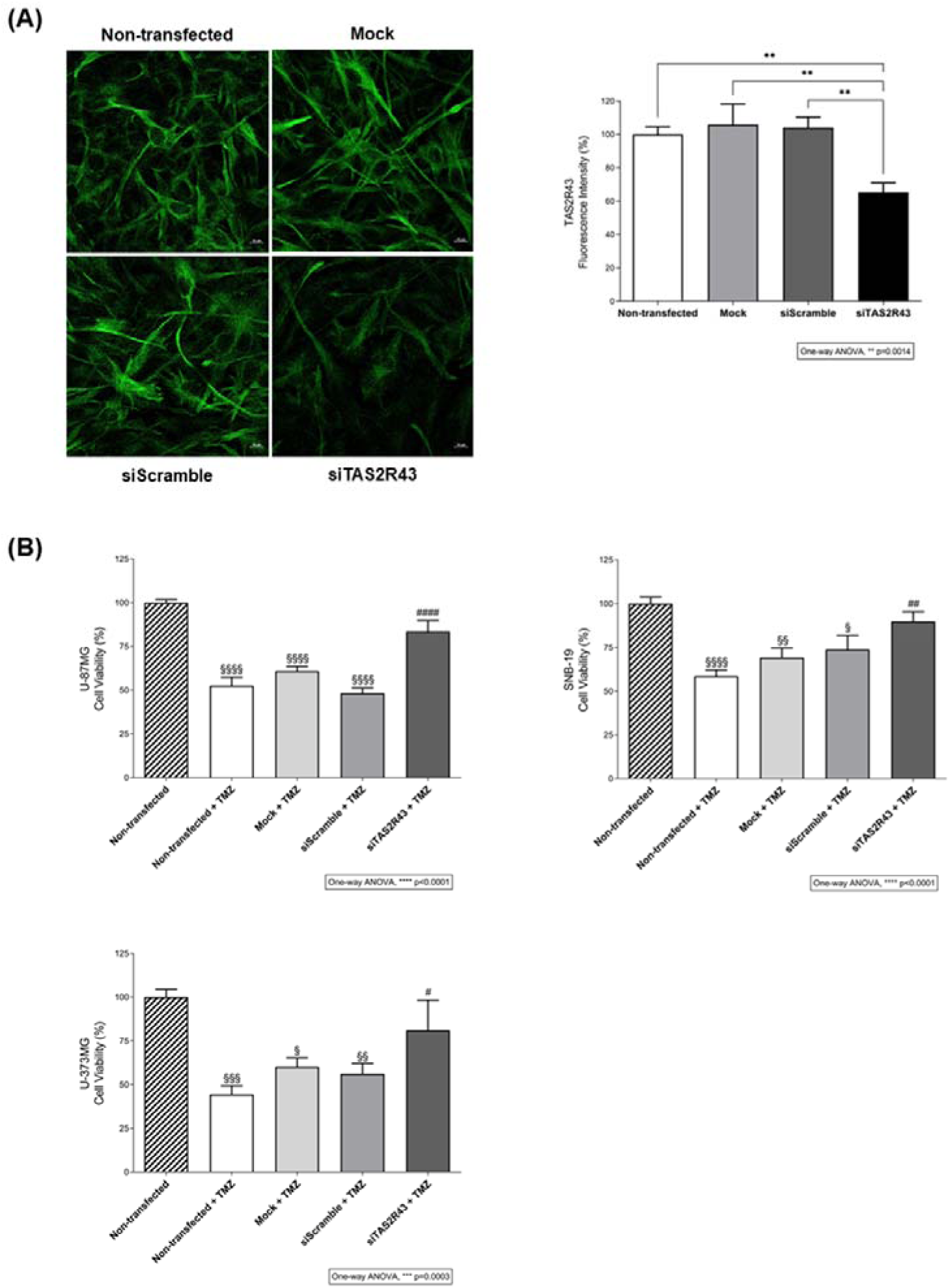
The anti-proliferative effects of temozolomide in glioblastoma cells depend on the activation of TAS2R43. (A) Immunofluorescence analysis of TAS2R43 expression after siRNA transfection in SNB-19 glioblastoma cells. Protein levels of siRNA TAS2R43-transfected cells are decreased in comparison with non-, mock-, and siRNA scramble-transfected cells. The quantification of TAS2R43 fluorescence intensity (green) was performed in different regions of interest of confocal microscopy images obtained from three independent experiments. Nuclei were stained with Hoechst 33342 (not shown). Scale bar: 10 µm. Bar graphs represent mean ± SEM. Statistical analysis was performed by one-way ANOVA followed by Tukey’s multiple comparisons test. [N=3 independent experiments; **p<0.01]. (B) Effects of TMZ in the viability of U-87MG, SNB-19, and U-373MG glioblastoma cells transfected or mock-transfected for 24 h with TAS2R43 or a scramble siRNA and incubated with 500 µM TMZ for 72 h. Bar graphs represent mean ± SEM. Statistical analysis was performed by one-way ANOVA followed by Tukey’s multiple comparisons test. [N=3 independent experiments; *p<0.05, **p<0.01, ***p<0.001 and ****p<0.0001; §versus non-transfected; #versus non-transfected+TMZ].

## 4. Discussion

The expression and functionality of the taste transduction pathway, particularly TAS2Rs, has been studied in several types of cancer, especially in breast and pancreatic cancer cells (Jeruzal-Świątecka et al. 2020), and associated with cancer chemoresistance. However, no studies have focused on glioblastoma or any other brain neoplasms despite its high levels of chemoresistance and limited treatment options. We addressed this gap by assessing the involvement of the taste signalling taste pathway in the response of human glioblastoma cells to TMZ, the golden standard treatment for these tumours, and predictably, a bitter taste compound. For these analyses, we chose three different glioblastoma cell lines with different morphology and proliferation rates: U-87MG are cluster invasion cells, whereas the most chemoresistant SNB-19 and U-373MG (similar to U-251MG) are individual invasion and expansive-growth cells, respectively (Diao et al. 2019; Memme et al. 2014).

In the last years, a growing body of evidence shows anti-proliferative, anti-invasiveness, pro-apoptotic, anti-angiogenic, and anti-metastasis effects of flavonoids, alkaloids, immunosuppressors, antibiotics, cannabinoids, and lactones in several types of cancer, including glioblastoma (reviewed in (Ammazzalorso et al. 2022; Costa et al. 2023; Luís et al. 2020)). Most of these compounds are known, or at least predicted, to be TAS2Rs ligands and some have also been reported as potential adjuvants in glioblastoma therapy with TMZ, like epigallocatechin gallate and resveratrol (Banerjee & Preissner 2018). The TASR14 and R39 ligand resveratrol has been extensively analysed in different cancers, including glioblastoma, where its ability to restrain cell proliferation, suppress tumour growth, and induce apoptosis in combination with TMZ was shown (Diogini et al. 2020; Li et al. 2016; Liu et al. 2020; Roland et al. 2013; Song et al. 2019; Yang et al. 2019). Also, the TAS2R39 ligand epigallocatechin-gallate was able to sensitize glioblastoma cells to TMZ and inhibit MGMT expression, decreasing cell viability and TMZ chemoresistance (Grube et al. 2018; McCubrey et al. 2017; Udroiu et al. 2019). Yet, the mechanisms underlying their anti-cancer effects mediated by TAS2R are still largely unknown, and the existing studies put in evidence varied signalling mechanisms. In a study carried out by Seo *et al*., in human neuroblastoma cells SH-SY5Y, TAS2R8 and R10 overexpression induced neurite elongation, decreased the expression of cancer stem cell markers, and inhibited self-renewal characteristics, as well as cell migration and invasion, and reduced tumour incidence and volume in mice (Seo et al. 2017). This suggests that these receptors could suppress the metastatic potential of neuroblastoma cells. Other studies in the human neuroblastoma cell line SH-SY5Y showed that probenecid inhibited salicin-induced ERK and CREB phosphorylation, the key transcription factor of neuronal differentiation, via TAS2R16 blockage, in this cell line (Wölfle et al. 2015). Others, showed that probenecid attenuated the HEK-293T cells’ response to the bitter compounds salicin, phenylthiocarbamide and 6-propyl-2-thiouracil, and aloin via TAS2R16, R38 and R43 inhibition, respectively (Greene et al. 2011). A recent study has highlighted the role TAS2R14 activation by lidocaine in head and neck squamous cell carcinoma cells, by inducing apoptosis through intracellular Ca^2+^ increase, mitochondrial dysfunction, and proteasome inhibition. This finding underscores the complex and varied signalling mechanisms involved in TAS2R-mediated anti-cancer effects and discloses novel therapeutic targets (Miller et al. 2023).

Although alkylating agents, like TMZ, are known for their strong bitterness and astringency, their potential to bind and activate specific TAS2Rs has not been addressed before. To ascertain the capacity of TMZ to activate the bitter taste signalling in glioblastoma cells, we performed functional assays with TMZ in the presence or absence of probenecid, a known TAS2R16, R38 and R43 antagonist (Greene et al. 2011; Wölfle et al. 2015), or by silencing the taste signalling pathway by GNAT3 knockdown. Our findings showed that TMZ reduced the cell viability, increased cell apoptosis, and elicited intracellular Ca^2+^ levels in a dose-dependent manner in U-87MG, SNB-19 and U-373MG glioblastoma cells. Interestingly, these TMZ effects were significantly reduced in the presence of probenecid, and GNAT3 knockdown, suggesting that the activation of the bitter taste signalling pathway potentiates somehow the anticancer effects of TMZ. The next step was the identification of the TAS2R to which TMZ binds. Based on *in-silico* analysis to predict the potential target-receptors for TMZ, TAS2R expression profiling in these cells, and probenecid binding, TAS2R43 emerged as the most likely TAS2R interacting with TMZ. In fact, when we silenced TAS2R43, the effects of TMZ in the glioblastoma cells analysed were also reduced similarly to what was seen with GNAT3 silencing and probenecid incubation. Thus, the expression of TAS2R43 in these cancer cells is essential for an effective action of TMZ. Interestingly, TAS2R43 is also a target of the bitter compound epigallocatechin gallate, which in turn has been widely used as an adjuvant agent in the treatment of gliomas, especially in glioblastoma, enhancing the therapeutic efficacy of TMZ (Chen et al. 2011; Grube et al. 2018; Xie et al. 2018; Zhang et al. 2015).

One of the major hurdles in cancer therapy is overcoming the action of efflux transporters to improve treatment efficacy. This is particularly troublesome for brain neoplasms since drugs have also to overcome the blood brain barrier where efflux transporters are also very active. Previous studies showed that TMZ is a substrate of MDR1 encoded by ABCB1, and we have previously demonstrated that TAS2R have the ability to control efflux transporters (Duarte AC, et al. 2020b; Munoz et al. 2015).

Since TMZ’s is well-known for its alkylating effects on DNA eliciting DNA damage and cell death, we propose that by binding TAS2R43, TMZ enhances its own penetrance into the cells possibly through the modulation of efflux transporters, as seen upon the activation of other TAS2Rs, while its blockage prevents its entrance to the cells. One of these transporters, the multidrug resistance protein 1 (MDR1), is expressed in low- and high-grade gliomas, and the intrinsic resistance of glioblastoma cells to anticancer drugs, including TMZ (De Faria et al. 2009; Gomez-Zepeda et al. 2019), has been attributed to this efflux transporter. This also explains the adjuvating effects of other TAS2R43 ligands like epigallocatechin gallate that enhance the anti-cancer effects of TMZ.

In summary, herein we demonstrated that TAS2R43 is activated by the chemotherapeutic alkylating drug TMZ, thereby contributing to its anti-proliferative and anti-apoptotic effects. These findings stress the importance of the taste transduction pathway via TAS2R43 suggesting that TAS2R43 could serve as a biomarker for the efficacy of temozolomide and other anti-cancer drugs that are substrates of MDR1.

## Acknowledgements

This work was supported by the Fundação para a Ciência e Tecnologia (FCT, Portugal) project grants (PTDC/BIM-ONC/7121/2014, UID/Multi/00709/2013 and UID/Multi/00709/2019), by CENTRO 2020 and Lisboa 2020 project grant (POCI-01-0145-FEDER-016822), and FEDER funds through the POCI – COMPETE 2020 – Operational Programme Competitiveness and Internationalization in Axis I – Strengthening research, technological development and innovation (POCI-01-0145-FEDER-007491). Ana R. Costa was recipient of a PhD fellowship (UI/BD/151025/2021) funded by FCT through the Portuguese state and EU budgets through the European Social Fund. Ana C. Duarte was recipient of a grant from CENTRO 2020 program through the ICON project (Interdisciplinary Challenges On Neurodegeneration; CENTRO-01-0145-FEDER-000013). We also acknowledge the support of the Portuguese Platform of Bioimaging (PPBI) [PPBI-POCI-01-0145-FEDER-022122] and the resources provided by the Fluorescence Microscopy Unit of RISE-Health, UBI.

